# A Taxonomic Revision of the Monocaul Phanerophyte *Ardisia* (Primulaceae) of Gabon

**DOI:** 10.64898/2026.06.26.734757

**Authors:** Harry Murdoch, Martin Cheek

## Abstract

Monocaul *Ardisia* (Primulaceae) have a single, vertical, woody stem and spiral phyllotaxy. They range from 30 cm to 100 cm tall. Three species of this architectural group, *Ardisia mayumbensis*, *A. hallei* and *A. bracteata,* have been recorded from Gabon hitherto. In this taxonomic revision we show that *A. mayumbensis* does not occur in Gabon, and we describe three new species. Two of these, *Ardisia doudou* sp. nov.and *A. mica* sp.nov., are endemic to Gabon and one, *A. litterbin sp.nov*, is found in both Gabon and Republic of the Congo. *Ardisia hallei* and *A. bracteata* are redescribed. All five species have a litter-gathering habit with a terminal funnel of leaves, and two of these species, *Ardisia doudou* and *Ardisia litterbin*, also possess adventitious roots in the distal part of the stem, a well-established strategy found in litter-gathering forest species of other plant families in tropical Africa. We provisionally assess the conservation status of all five taxa using the 2012 IUCN standard, finding that all monocaul *Ardisia* in Gabon fall within threatened categories. Two of the species, *Ardisia bracteata* and *A. mica,* are known from single collections and have not been seen for 164 and 63 years respectively and are conceivably extinct although further surveys are needed to establish this. We employ new characters in delineating and describing African *Ardisia* using leaf thickness and oil gland data.

## Introduction

*Ardisia* Sw. (Swartz 1788) is a pantropical genus in the family Primulaceae (subfamily Myrsinoideae) including 739 accepted species (POWO continuously updated). The genus is distributed in the Americas, Asia and Africa. African *Ardisia* face several taxonomic challenges including the loss of 17 holotypes in the 1943 bombing of the Berlin Herbarium, many of which lack isotypes (Taton 1979).

The first two *Ardisia* described from Africa were *A. cymosa* Baker from São Tomé and *A. bracteata* Baker from Gabon (Baker in Oliver 1877). In 1902, eight species from Cameroon were described (Gilg 1902). Mez (1902) erected a new genus, *Afrardisia* Mez, for the ten then known African *Ardisia*. A further seven species of *Afrardisia* were described, mainly endemic to Cameroon, one from both Cameroon and DRC, and one endemic to DRC (Gilg and Schellenberg 1913). In the following four decades, three more *Afrardisia* were described, including one from DRC (De Wildeman and Bequaert 1925) and *Afrardisia mayumbensis* Good from Angola (Cabinda) (Good 1927). De Wit’s revision of *Afrardisia* (1958) synonymised six species and named a further new species, *Afrardisia comosa* de Wit.

Taton (1979) demonstrated that the basis for separating *Afrardisia* from *Ardisia* (ovule placentation), as used by Mez (1902) and followed by De Wit (1958), were not upheld, and transferred all *Afrardisia* into *Ardisia*. In this revision, he made six new combinations from *Afrardisia*, as well as publishing two new names, and 17 new African *Ardisia* species. Among these are 10 species considered by Taton to be from Gabon: *A. buesguenii* (Gilg & Schellenb.) Taton, *A. mildbraedii* (Gilg & Schellenb.) Taton, *A. mayumbensis* (R. Good) Taton, *A. atrobullata* Taton, *A. bampsiana* Taton, *A. lethomasiae* Taton, *A. pierreana* Taton, *A. hallei* Taton, *A. letestui* Taton and *A. belingaensis* Taton. In addition, Taton cited specimens from Gabon for what he treated as *A. sadebeckiana* Gilg and *A. staudtii* Gilg, both previously considered to be endemic to Cameroon. Taton included short descriptions of five unnamed species (‘sp 1’ to ‘sp 5’) because material was limited. In this paper we formally describe one of these species to improve the possibility of it being protected from extinction.

Cheek (in Cheek et al. 2010) published two new species from Cameroon. Cheek and Xanthos (2012) described *A. ebo* Cheek from Cameroon and Gabon an herbaceous species. Peng and Cheek (2025) accomodated nine species with a herbaceous, creeping habit in a newly erected subgenus, *Ardisia* subg. *Kamardisia* Cheek. These included six new species, five of these are Gabon endemics, and one is known from both Cameroon and Gabon (Peng and Cheek 2025).

At the time of writing, 51 species of *Ardisia* are recognised from continental Africa, Sixteen species are recognised in the Gabon checklist (Sosef *et al*. 2006) and the further six in Peng and Cheek bring the total to Gabon to 22 species. This paper continues the effort towards a Flore Du Gabon account for *Ardisia*.

African *Ardisia* species have been classified into four groups based on lifeform (Cheek et al. 2025). In this paper we focus on group 3, the ‘monocaul phanerophyte’ *Ardisia* of Gabon. These have a single, woody, vertical stem with spiral phyllotaxy, and lack axillary branches. They range from 30 cm to 100 cm tall. They are distinct from both the group 4 ‘branching phanerophyte’ *Ardisia*, which have horizontal branches with distichous phyllotaxy arising from a vertical stem with spiral phyllotaxy, and range from 0.5 m to 20 m tall, and from the group 2 ‘erect chamaephytes’ (herbaceous with erect single stems with distichous phyllotaxy) and group 1, creeping herbaceous *Ardisia* which do not exceed 20 cm in height (Peng and Cheek 2025). The monocaul *Ardisia* currently accepted in Gabon are *A. mayumbensis*, *A. hallei* and *A. bracteata* (Sosef et al. 2006). Here we revise the monocaul phanerophyte *Ardisia* of Gabon. We describe three new monocaul *Ardisia* endemic to Gabon and show that *A. mayumbensis*, long considered a Gabonese species, does not occur in Gabon. *A. hallei* and *A. bracteata* are given new, more detailed descriptions.

Of the three currently accepted monocaul *Ardisia* from Gabon, only one (*A. hallei*), has received any kind of conservation assessment (Paradis et al. 2024). Here we provide provisional conservation assessments (using IUCN 2012 criteria) for the four other monocaul species of Gabon. Many species of African *Ardisia* are range-restricted and threatened by clearance or disturbance of the evergreen forest habitat to which they are all restricted. In neighbouring Cameroon, 16 species are listed as threatened in the Red Data Book (Onana & Cheek 2011).

Multiple recent studies have found pantropical *Ardisia* not to be monophyletic (Julius et al. 2021). Larson *et al*. (2023) find at least 19 smaller genera nested within *Ardisia*. Yang and Hu (2022) describe a genus-level taxonomic revision as “necessary”. Considering this and given that the type species of the genus is *A. tinifolia* Sw. from Jamaica, it is conceivable that *Ardisia* will be split up and that African *Ardisia* will take on a distinct name (or names) in the future, so that *Afrardisia* Mez may need to be restored.

Many of the specimens treated in this paper have previously been determined as *A. mayumbensis*, and the working name for this paper was the “*A. mayumbensis* complex in Gabon”. However, having delimited the taxa, we found that *A. mayumbensis* itself does not occur in Gabon, but in Republic of the Congo, DRC and Angola (Cabinda). We found that Taton (1979) had included a specimen, *Chevalier* 27044 (P) from Gabon within his concept of *A. mayumbensis*, but with the benefit of many more specimens it is clear that this specimen is discordant with the type of *A. mayumbensis* from Angola. All *A. mayumbensis* specimens cited in Sosef *et al*.’s checklist (2006) have here been accounted for and placed either as *A. doudou* or *A. litterbin*. The differences between these three species are given in Table 1. In addition the two new species are diagnosed against *A. mayumbensis*. Only one specimen cited by Taton (1979) and also by Sosef et al. (2006) as *A. mayumbensis* is not treated here. This is *N. Hallé* 1330, which clearly demonstrates a branching habit, and is not a monocaul.

It is quite possible that with further collection efforts *A. mayumbensis* could yet be found in Gabon because the closest specimen to Gabon of that species, *Nkondi* 673, is found in Republic of the Congo only c. 14 km from the border with Gabon. Additional monocaul species occur in Cameroon but are beyond the scope of this paper. We plan to address these in a further paper. None of the Gabonese monocaul species occur in Cameroon or vice versa.

## Material and methods

This taxonomic treatment was undertaken entirely with herbarium specimens. Specimens seen in person include those from Royal Botanic Gardens, Kew (K) as well as loans from the National Herbarium of the Netherlands, Wageningen (WAG). Visits were made to P and BM in July 2025 to see specimens there. Field notes have been vital here in identifying characters lost in the drying and pressing process. Protologues have been referred to throughout.

Measurements and descriptions were made directly from herbarium specimens. All microscopic measurements of specimens seen at K were performed with a Leica S9E dissecting microscope. Descriptions and measurements of leaf scales were made with a Leica M165C. Specimens seen at P were examined and measured with an equivalent Nikon stereo microscope. Leaf thickness measurements were made with a Digital Micrometers Ltd DML3033 Digital Micrometer which measures to 0.01mm digital thickness. Leaf blade backlighting has been undertaken with an iPhone 13 torch placed under the specimen facing up and viewed through a dissection microscope. Terminology for the leaf descriptions follows the Leaf Architecture Working Group (1999). Genus-specific terminology mostly follows Taton (1979), making reference also to Cheek and Xanthos (2012) and Peng and Cheek (2025). Description of all other morphology follows Beentje & Cheek (2003). All specimens were seen unless indicated as “not seen”.

Important in separating species here has been the size, shape and quantity of oil glands, which are found in the leaves and floral parts of all African *Ardisia* seen. Of particular importance are the oil glands in the leaf lamina, the density, also the shape and size of which, as can be seen in Table. 1, varies considerably between species but can be remarkably consistent within species. Using oil glands as characters for species delimitation was established by Peng and Cheek (2025) and has been developed as part of this paper.

The species distribution map (Fig. 1) was created in R (R Core Team 2025) using the following packages: ggplot2 (Wickham 2016), dplyr (Wickham et al. 2023), sf (Pebesma 2018, Pebesma and Bivand 2023), rnaturalearth (Massicotte and South 2025), rnaturalearthdata (South et al. 2024) and readr (Wickham et al. 2024). Protected area data is from The World Database on Protected Areas (UNEP-WCMC and IUCN 2025).

**Fig. 1.**
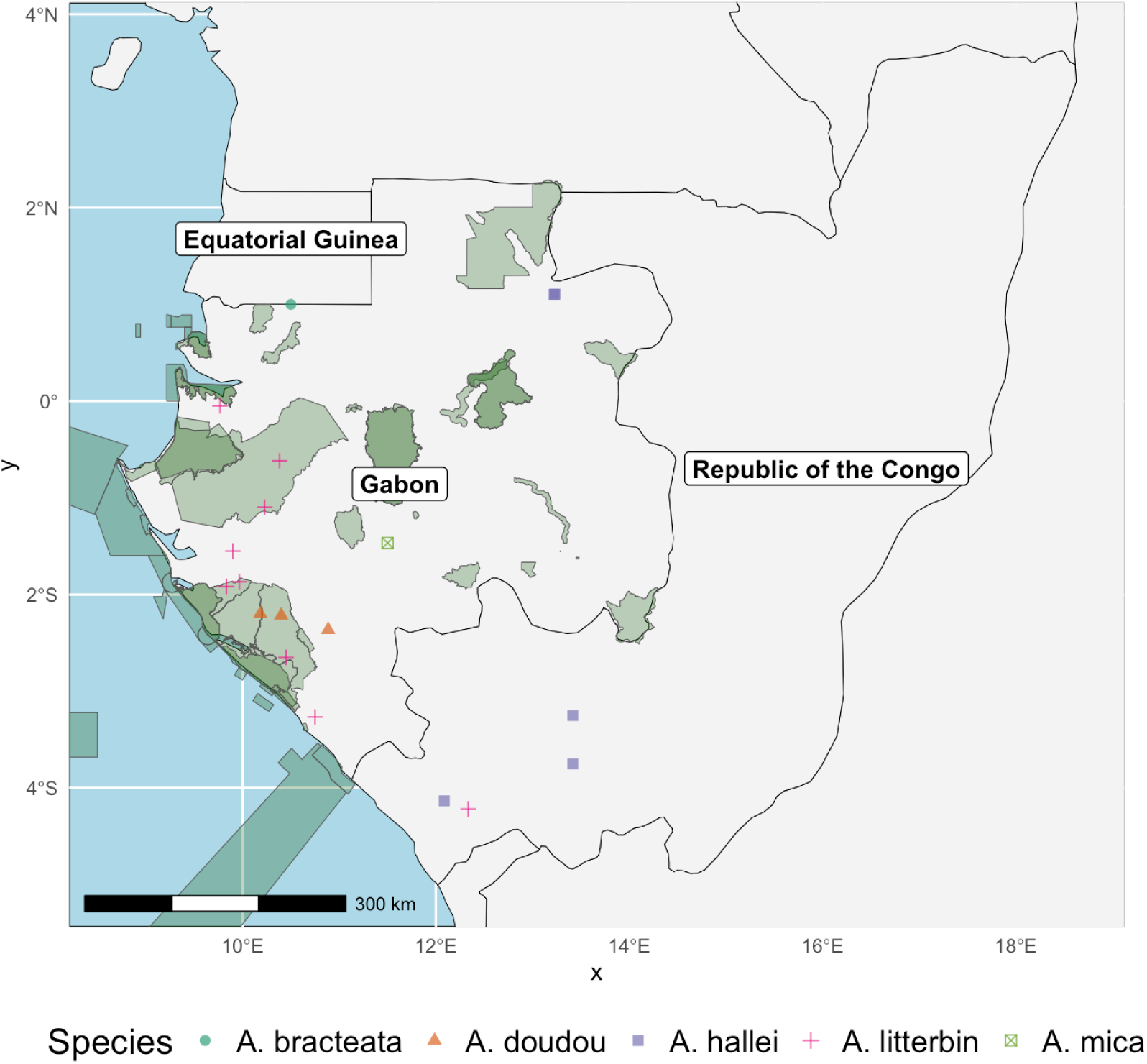
A global species distribution map of the five monocaul *Ardisia* species of Gabon, including two species (*A. hallei* and *A. litterbin*) which extend to Republic of the Congo. Protected areas overlayed in green. Note that the most northerly point for *A. hallei* represents two specimens collected on different occasions with very similar estimated coordinates.

To assess conservation status and for a distribution estimate, specimens without coordinates were georeferenced using Google Earth Pro (continuously updated) based on information on specimen labels. Conservation status was assessed following the IUCN Red List categories and criteria (IUCN 2012; IUCN Standards and Petitions Committee 2024). The Area of Occupancy (AOO) and Extent of Occurrence (EOO) were calculated using GeoCAT (Bachman et al. 2011). The AOO was calculated using a default cell size of 4 km². Global Forest Watch (World Resources Institute 2014) is used to assess tree cover loss or gain within a species range, as well as to view logging concessions and oil palm plantations. Google Earth satellite imagery is used to check locations for deforestation. International and local news sources are cited where relevant.

## Taxonomic treatment

### 1. *Ardisia doudou* Cheek & Murdoch sp. nov. (Fig. 2)

Syn. *Ardisia mayumbensis* sensu Sosef *et al*. (2006) pro parte quoad. Sosef et al. 1226, J.J.F.E. de Wilde et al. 8988, non (R. Good) Taton (1979)

**Fig. 2.**
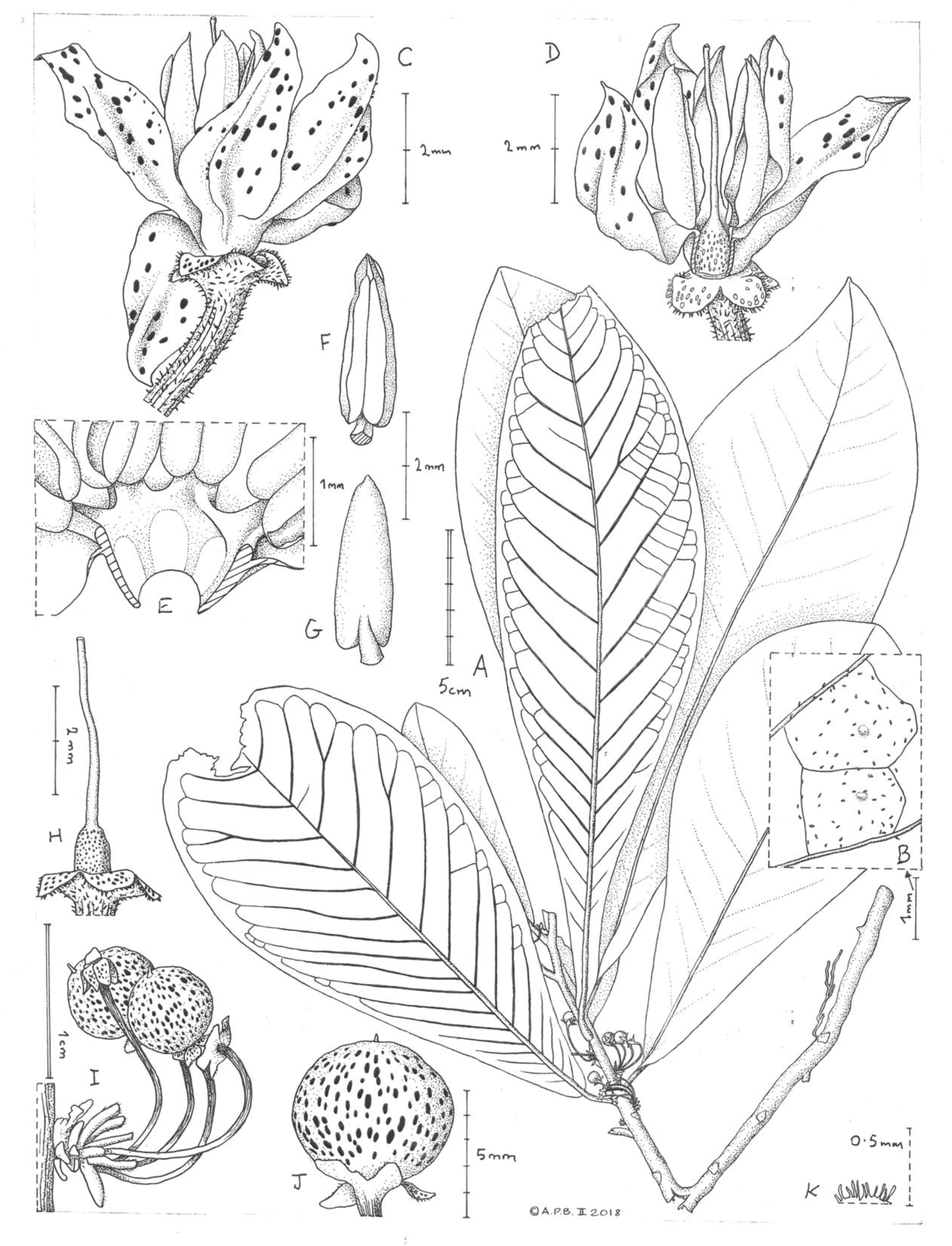
Ardisia doudou. A habit (illustration is of atypical specimen without dense adventitious roots); B details of leaf abaxial surface showing areolae, oil glands and 1-3-armed scales; C flower, lateral view (petal at rear recurved); D as C with petal and one stamen removed; E dissection of fused corolla and androecial tube; F anther, ventral face; G anther, dorsal face; H ovary, style and stigma in situ; I inflorescence in fruit; J fruit; K detail of fimbriate margin of calyx lobe. A, B & J *J.J.F.E. de Wilde & Jongkind* 9527, C-H *J.J.F.E. de Wilde et al.* 8988. (A adventitious roots added from *J.J.F.E. de Wilde et al.* 8988.) All drawn by Andrew Brown.

#### Diagnosis

*Ardisia doudou* is similar to *Ardisia litterbin* Cheek & Murdoch and *A. mayumbensis*, differing in the pedicel posture ascending in fruit (vs descending in *A. litterbin* and *A. mayumbensis*), gland density (leaf blade, no. per 2 × 2 mm) 2–6 (vs *A. litterbin* 8–16 and *A. mayumbensis* 10–27), leaf thickness (lamina) 0.7–0.75 mm (vs *A. litterbin* 0.31–0.53 mm and *A. mayumbensis* 0.25–0.31 mm). The anthers lack surface oil glands (unique among all Gabonese phanerophyte monocaul *Ardisia* species). See Table 1 for more diagnostic characters.

### Type

Gabon, Ogooue-Maritime, Doussala, Projet WWF sur la Biodiversité des Monts Doudou, a plus ou mois 40 km au Nord-Ouest de Doussala, autour du campement II. Le long d’une riviere, 2°13’S, 10°24’E, 450 m, fr. 11 April 2000, Sosef et al. 1226 (holotype K; isotypes WAG, P, MO, not seen).

#### Etymology

Named as a noun in apposition for the Doudou Mountains, from where all known specimens were collected.

#### Description

Monocaul, 50–60 cm tall, stem vertical, unbranched, ground roots not seen, phyllotaxy spiral, terete, 5–11 mm diam., internodes 3–18 mm long, rusty brown tomentellous indumentum on the distal 3–4 nodes, glabrescent, epidermis red brown to grey brown, darker towards apex; Adventitious roots usually present at the distal 5–10 nodes, roots 8–26 mm long, often densely branched and matted with organic matter, root hairs glabrescent to rusty brown floccose, often dense distally, becoming sparse proximally. Leaves clustered towards stem apex, 5–6 leaves in the distal 5.5–7.5 cm of stem. Petiole canaliculate, 4–13 × 3–5 mm; lamina discolorous, adaxially dark green and dull in life, abaxially paler green in life, coriaceous in life,oblanceolate, 19.2–33.3 × 8.6–14 cm, length to width ratio 2.2–3.4:1, apex acute to rounded, base gradually attenuate to cuneate, margin entire, lamina c. 0.7 mm thick, c. 1.6 mm at midrib, midrib conspicuous, darker green than lamina in life, raised on the adaxial and abaxial surfaces, grooved on adaxial; lateral nerves 14–29 on each side of the midrib, departing from midrib at 40–60 degrees, proximal 1/3^rd^ to 2/3^rd^ eucamptodromous, distal 1/3^rd^ to 2/3^rd^ brochidodromous, looping 3–8 mm from margin, tertiary nerves conspicuous, raised abaxially, reticulate to scalariform creating areolae; oil glands black to orange (transmitted light), brown to black in reflected light, inconspicuous abaxially (in reflected light), visible and raised on adaxial surface in reflected light, densest near margin, subcircular, elliptic or oblong, 0.15–1.27 × 0.11–0.2 mm, 2–6 per 2 mm × 2 mm; 1–4(–6) glands per areole; scales conspicuous (microscope), 1–3(–6)-armed, reddish brown, on abaxial and adaxial surfaces, 14–22 per 2 mm × 2 mm abaxially, 1–7 per 2 mm × 2 mm adaxially, 0.03–0.13 mm wide. Inflorescences cauliflorous, 0–2 per internode, 1–7-flowered, often among adventitious roots, peduncle usually inconspicuous or up to 3 mm long, rachis 4–6 mm long, pedicel bases exserted from the rachis, rusty brown floccose hairs on rachides and bracts; bracts triangular-lanceolate, 2–5 × 0.7–1.2 mm; pedicels ascending, u-shaped, terete, 10–25 × 0.5–1 mm, rusty brown pilose and dark reddish brown oil glands at surface throughout, oil glands raised, subcircular, elliptic or oblong, 0.05–0.2 × 0.05–0.2 mm. Flower buds (J.J.F.E. de Wilde et al. 8988) narrowly ovoid, c. 5 × c. 2 mm, apex acute; calyx appressed to reflexed, lobes 5, overlapping to the right, ovate, c. 0.8 × c. 0.9 mm, apex acute to rounded, oil glands covering 30% of calyx, orange to brown, slightly raised on inner surface, subcircular, elliptic or oblong, c. 0.06 × c. 0.06 mm, margin fimbriate, fimbriae translucent, 0.05 – 0.075 mm long, corolla twisted slightly to the right. Flowers (J.J.F.E. de Wilde et al. 8988) 5-merous, calyx as bud. Corolla spreading, off-white with pale red dots (likely oil glands) in life, c. 7 mm diam., tube shortly cylindric, c. 1 × c. 1.1 mm staminal tube adnate ; corolla lobes narrowly ovate, 2.6–3.5 × 1.1–1.4 mm, apex acute, glands sparse, c. 10% of surface, visible on both sides, orange, subcircular, elliptic or oblong, 0.13–0.22 × 0.07–0.18 mm. Androecium erect, staminal tube shortly cylindric, c. 0.7 × 1.1 mm, filaments dorsiventrally flattened, c. 0.7 × 0.4 mm, widest at base; anthers narrowly elliptic, 2.3–2.6 × 0.8–0.9 mm, base subsagittate, apex acute, subbasifixed, oil glands absent. Ovary subcylindric, 0.8–1 × 0.7 mm, oil glands raised, covering c. 40% of the distal surface, dark reddish brown, subcircular, elliptic or oblong, 0.03–0.07 × 0.03–0.07 mm, style terete, c. 3 mm long, stigma not seen. Fruits orange to red in life, subglobose, 6–7 mm diam., calyx reflexed, otherwise as bud, bracts persistent, style base persistent, c. 0.3 × c. 0.24 mm; oil glands covering c. 30% of surface, less dense around base, dark reddish brown, elliptic to lanceolate, 0.2–0.5 × 0.1–0.3 mm, longest near calyx, glabrous, ovules c. 5, irregularly arranged; fruit wall c. 0.02 mm thick.

### Phenology

Flowering November; fruiting March and April.

### Distribution and ecology

Doudou Mountains, Gabon. 170–450 m alt. Occurs as an understory rainforest shrub. Adventitious roots, which are matted with organic matter, indicate an adaptation to collect falling litter as a means of gathering nutrients.

### Conservation status

*Ardisia doudou* is known from only three specimens within or close to the Doudou Mountains in southwest Gabon, representing two locations. The species has an Extent of Occurrence (EOO) of 149 km² and Area of Occupancy (AOO) of 12 km². Two specimens (Sosef et al. 1226 and J.J.F.E. de Wilde et al. 8988) were collected within the protected area of the Moukalaba-Doudou National Park (UNEP-WCMC and IUCN 2025). This area was already protected when the collections were made (1986–2000), though the area had seen extensive logging prior to this. Global Forest Watch shows almost no tree cover loss between 2001 and 2024 in this protected area (World Resources Institute. 2014). Satellite imagery (viewed July 2025, (Google Earth n.d.)) shows intact forest at both collection sites with no immediate threats.

J.J.F.E. de Wilde & Jongkind 9527 was collected just outside of the national park and is not from a protected area. This location occurs within a logging concession (World Resources Institute. 2014) and is under 35 km from an oil palm plantation to the east. The town of Moabi is 10 km to the South and includes a small airport. Satellite imagery shows some forest clearance around the town, with at least some of this being clearly used for agriculture. The possible expansion of either this town or oil palm plantations nearby represent a possible future threat to this site Within the AOO of all three collections, Global Forest Watch shows < 0.1% tree cover loss between 2000 and 2024 (World Resources Institute, 2014). Ecotourism activities take place in the area, though are mainly focused on animals including gorillas and chimpanzees. Outside of protected areas, we infer a current and a future continuous decline in the extent and the quality of its habitat due primarily to logging, as well as the possible expansion of oil palm plantation and urban areas. Due to these threats, as well as the restricted EOO and AOO, here we assess this species provisionally as Endangered B1ab(iii)+ B2ab(iii). **Notes.** *A. doudou* is unique among all known African monocaul species in the pedicels, which curve upwards, positioning the fruit towards the stem apex (Fig.1). Also unique is the lack of surface oil glands on the anthers. One of only two African *Ardisia* (alongside *A. litterbin*) to collect organic matter in adventitious roots in the apical part of the stem. All flower measurements were made from three already dissected flowers in packets, from two sheets of one collection (J.J.F.E. de Wilde et al. 8988).

### Additional specimens examined (paratypes)

Gabon. Ogooue Maritime, Doussala, Projet WWF sur la Biodiversité des Monts Doudou, a plus ou mois 40 km au Nord-Ouest de Doussala, autour du campement II. Le long d’une riviere, 2°13’S, 10°24’E, 450 m, fr. 11 April 2000, Sosef et al. 1226 (holotype K; isotypes WAG, P, MO, not seen); Doudou Mountains, about 60 km along exploitation track in WNW direction from Doussala, exploited high forest, 2°12’S, 10°11’E, 200 m, b., fl., 27 Nov. 1986, J.J.F.E. de Wilde et al. 8988 (WAG); Doudou Mts, about 88 km NNW of Doussala, on the track towards Bongo. Closed wet high forest, 2°22’S, 10°52’E, 150 m, fr., 23 March. 1988, J.J.F.E. de Wilde & Jongkind 9527 (WAG)

### 2. *Ardisia litterbin* Cheek & Murdoch sp. nov. (Fig. 3)

Syn. *Ardisia mayumbensis* sensu Sosef *et al*. (2006) pro parte quoad F.J. Breteler et al. 9480, C.C.H. Jongkind et al. 5746, van Nek 355, J.J. Wieringa 909, J.J.F.E. de Wilde 9203 non (R. Good) Taton (1979)

**Fig. 3.**
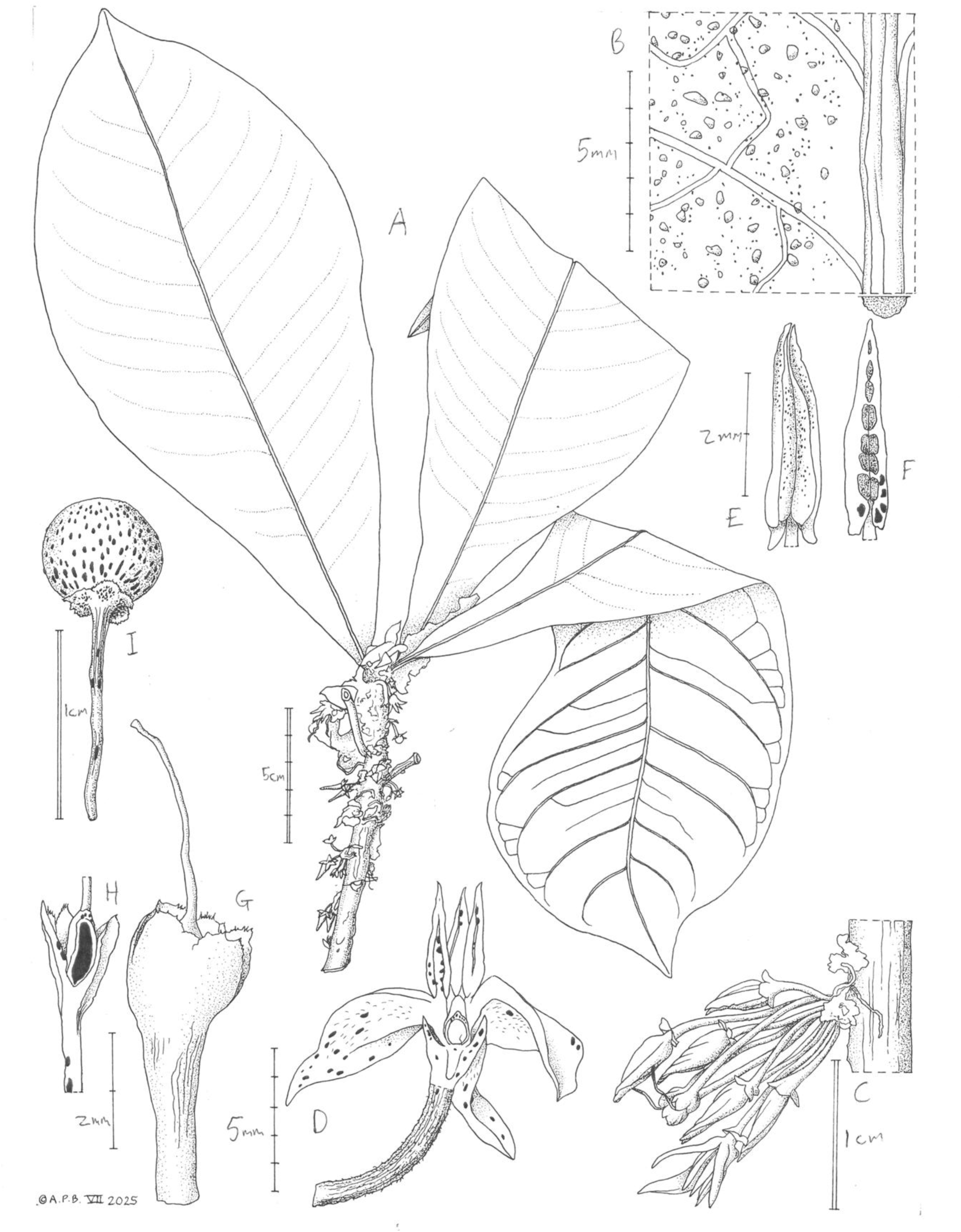
Ardisia litterbin. A habit; B leaf abaxial at midrib showing glands and scales (with profile of midrib); C inflorescence at bud stage; D half-flower (hydrated); E stamen (ventral face, dry); F stamen (dorsal face, dry); G ovary and style in calyx (dry); H cross section of ovary in calyx (specimen flattened); I mature fruit A, B *J.J.F.E. de Wilde et al.* 9203 sheet 1, C *J.J.F.E. de Wilde et al.* 9203 sheet 2, D-H *Jongkind et al.* 5746, I *F.J. Breteler et al.* 9480. All drawn by Andrew Brown.

Syn. *Ardisia mayumbensis* sensu (R. Good) Taton (1979) pro parte quoad Chevalier 27044

#### Diagnosis

*Ardisia litterbin* is most similar to *A. doudou* and *A. mayumbensis*, differing in gland density (8–16 per 2 × 2 mm of leaf blade vs *A. doudou* 2–6 and *A. mayumbensis* 10–27); glands per areole (leaf blade reticulation) 5–35 vs *A. doudou* 1–4(–6) and *A. mayumbensis* 4–55; leaf thickness (lamina) 0.31–0.53 mm vs *A. doudou* 0.7–0.75 mm and *A. mayumbensis* 0.25–0.31 mm; stem diameter (at 10 cm above ground) 6–8 mm vs *A. doudou* 5–11 mm and *A. mayumbensis* 3.5–5 mm, and longest leaf length 23.2–30 cm vs *A. doudou* 28.3–33.3 cm and *A. mayumbensis* 18–22 cm). See Table 1 for more diagnostic characters.

#### Type

Gabon, Nyanga, 22 km along the road from Mayumba to Tchibanga and then about 10 km along a forest exploitation track in an eastern direction. Exploited high forest near the Doussa river, 3°16’S, 10°45’E, +-10 m, b., fl. 07 Dec. 1986, J.J.F.E. de Wilde et al. 9203 (holotype WAG; isotypes all not seen: LBV, BR, MO).

### Etymology

Named as a noun in apposition for the plant’s litterbin habit. The “Litterbin” phrase is taken from field notes of Jongkind et al. 5746. The name means a receptacle for rubbish, referring to the field observations that this species collects fallen debris in its funnel of leaves.

### Description

Monocaul, 50–70 cm tall, stem vertical, unbranched, phyllotaxy spiral, terete, 6–8 mm diam., internodes 3–11 mm long, rusty brown tomentellous indumentum in the distal 3–5 nodes, becoming glabrescent, epidermis grey brown, darker towards apex; Adventitious roots usually present within the distal 5–18 nodes, 10–20 mm long, densely branched and often matted with organic matter, invertebrates visible; root hairs inconspicuous; Leaves clustered towards stem apex, 5–12 leaves in the distal 4–14 cm of stem. Petiole canaliculate, 5–16 × 2–3 mm; lamina discolorous, adaxially green and dull to glossy in life, abaxially paler green in life, young leaves purplish in life, coriaceous in life (J.J.F.E. de Wilde et al. 9203, F.J. Breteler et al. 9480, I. van Nek 355, J.J. Wieringa 909, J.J.F.E. de Wilde et al. 9887), oblanceolate, 17–30 × 6.4–12 cm, length to width ratio 2.4–3.4:1, apex attenuate to rounded, base gradually attenuate to cuneate, margin entire, lamina 0.4–0.5 mm thick, 0.8–1.5 mm thick at midrib, midrib conspicuous, raised on the adaxial and abaxial surfaces, grooved on adaxial; lateral nerves 14–20 on each side of the midrib, departing from midrib at 40–60 degrees, proximal 1/5^th^ to 2/5^th^ eucamptodromous, distal 3/5^th^ to 4/5^th^ brochidodromous, looping 2–6 mm from margin, tertiary nerves conspicuous, raised abaxially, reticulate creating areolae; oil glands black to orange-red (transmitted light), brown to black in reflected light, visible and raised on adaxial and abaxial surface in reflected light (lens needed), densest near margin, subcircular, elliptic or oblong, 0.11–1.45 × 0.11–0.27 mm, 8–16 per 2 mm × 2 mm; 5–35 glands per areole; scales conspicuous (microscope), 2–4-armed, reddish brown, on abaxial and adaxial surfaces, 14–23 per 2 mm × 2 mm abaxially, 0–10 per 2 mm × 2 mm adaxially, 0.03–0.13 mm wide. Inflorescences subaxillary or cauliflorous, 0–2 per internode, 1–15 flowered, often among adventitious roots, sessile, rachides 1.6–5 mm long, pedicel bases exserted from the rachis, rusty brown floccose hairs on rachides, bracts and pedicels; bracts caducous, ovate-lanceolate, 2–6 × 0.6–3 mm, dark reddish brown oil glands at surface, subcircular, elliptic or oblong, 0.11–0.27 × 0.11–0.18 mm; pedicels descending, terete, 3–14 × 0.5–1 mm, oil glands variable, range from inconspicuous to as bracts. Flower buds ovoid to narrowly ovoid, 5–7 × 2–4 mm, apex narrowly acute, calyx cupular to reflexed, lobes 5, overlapping to the right, ovate, 0.7–2 mm × 0.8–1.8 mm, apex acute, oil glands covering 30% of calyx, dark orange brown, slightly raised on outer surface, subcircular, elliptic or oblong, 0.05–0.29 × 0.05–0.15 mm, margin fimbriate, fimbriae translucent, 0.05–0.1 mm long, corolla twisted to the right. Flowers 5-merous, calyx as bud. Corolla spreading to reflexed, pale purple in life (I. van Nek 355, J.J.F.E. de Wilde et al. 9887, C.C.H. Jongkind et al. 5764), 8–12 mm diam., tube shortly cylindric, c. 0.9 × c. 0.8 mm adnate to staminal tube; corolla lobes narrowly ovate, 3.9–5.5 × 0.9–2 mm, apex acute, glands sparse, c. 10% of surface, visible on both sides, orange, subcircular, elliptic or oblong, 0.11–0.27 × 0.07–0.18 mm. Androecium erect, staminal tube shortly cylindric, c. 1.4 × 0.8 mm, filaments dorsiventrally flattened, c. 0.3 × 0.2 mm, widest at base; anthers narrowly elliptic, 3.5–4 × 0.9–1.3 mm, base sagittate, apex acute, subbasifixed, oil glands covering c. 20% of the dorsal surface scattered over both anther cells, conspicuous, dark brown to black, subcircular, elliptic or oblong, 0.07–0.2 × 0.07–0.1 mm, dissipating towards the margin, ventrally inconspicuous or absent. Ovary subglobose, c. 1.8 mm diam., style terete, c. 5 mm long, stigma c. 0.2 mm. Fruits dark orange in life, subovoid to subglobose, 7–8.5 mm diam., calyx flat to reflexed, otherwise as bud, style base persistent or caducous, c. 0.6 × c. 0.2 mm; oil glands covering c. 25% of surface, black to dark reddish brown, subcircular, elliptic or oblong, 0.2–0.8 × 0.1–0.2 mm, longest near calyx; glabrous, ovules c. 5 (F. J. Breteler et al. 9480), irregularly arranged; fruit wall c. 0.07 mm thick.

### Phenology

Flowering May, October to December, fruiting March, May and November.

### Distribution and ecology

Gabon and Republic of the Congo. Along coastal mountains from the Crystal Mountains in Northern Gabon to the Mayombe Forest in Southern Gabon and Republic of the Congo. Understory rainforest shrub occurring at 10 – 460 m elevation. Adventitious roots, which are matted with organic matter, indicate an adaptation to collect falling litter as a means of gathering nutrients.

### Conservation status

Known from a total of nine collections across seven locations. Eight collections span the length of the coast of Gabon and one collection is from western Republic of the Congo. The species has an EOO of 42,652 km² and an AOO of 36 km². In Gabon, six collections come from protected areas, with the remaining two unprotected. The one collection from Republic of the Congo comes from a nominally protected area (Reserve de la Biosphere de Dimonika). Despite being in a protected area, F.J. Breteler et al. 9480 is within a kilometre of an oil exploration and production operation by Assala Energy, acquired by Gabon’s national oil company in 2024 having increased its production by 30% since 2017 (Mishra 2024). Similarly, J.J. Wieringa 909 is in a protected area but is less than three kilometres from a logging concession, which crosses into the protected area. Global Forest Watch shows a 9.9% tree cover loss within a 2 km cell size around this collection site between 2000 and 2024 (World Resources Institute. 2014).

J.J.F.E. de Wilde et al. 9887 occurs outside of a protected area and, while its precise location is unclear, is likely to occur within or close to a logging concession. The J.J.F.E. de Wilde et al. 9203 collection site is not in a protected area and is found within a logging concession. Despite most collections coming from nominally protected areas, there are clear threats to this species, primarily from logging, and also from oil extraction. From this we can infer a continued decline in area, extent and quality of habitat. Due to these threats, as well as its restricted AOO, here we provisionally assess this species as Vulnerable B2ab(iii).

### Notes

One of only two African *Ardisia* (alongside *A. doudou*) to collect organic matter in a funnel of leaves, together with adjacent adventitious roots in the apical part of the stem. Litter also collects in the longitudinal groove of petioles. Flowers are positioned below the adventitious roots, which may a strategy to improve access for pollinators

#### Additional specimens examined (paratypes)

Gabon. Aledjom sur le Remboue, 0°03’N, 09°46’E, 12 Oct. 1912, Chevalier 27044 (P); Wolen-Ntem. WNW of Tchimbele. Primary forest, 0°37’S, 10°23’E, 450 m, fr., 12 May 1990, J.J. Wieringa 909 (WAG); Moyen-Ogooue, Zone de Lambaréné, à 50 km au sud de la ville, à 15 km au sud-est du lac Ezanga. Forêt des bas-fonds humides, 01°05’49’S, 10°13’46’E, 48 m, fr., 20 Nov. 2013, Bidault et al. 1456 (P); Rabi 12. More or less primary dryland forest, 1°33’S, 9°54’E, 50 m, b., fl., 30 Nov. 1989, J.J.F.E. de Wilde et al. 9887 (WAG); Forest near Checkpoint, 1°52’S, 9°58’E, b., fl., 17 Nov. 1990, I. van Nek 355 (WAG); Rabi. In rainforest, 1°55’S, 9°50’E, fr., 25 March 1990, F.J. Breteler et al. 9480 (WAG) Nyanga, Doudou Mountains, Chantier SFN-Bakker. Valley with small stream in forest, 2°39.1’S, 10°27.0’E, 200 m, b., fl., 22 Nov. 2003, C.C.H. Jongkind et al. 5746 (WAG); Nyanga, 22 km along the road from Mayumba to Tchibanga and then about 10 km along a forest exploitation track in an eastern direction. Exploited high forest near the Doussa river, 3°16’S, 10°45’E, +-10 m, b., fl. 07 Dec. 1986, J.J.F.E. de Wilde et al. 9203 (holotype WAG; isotypes LBV, BR, MO). Republic of the Congo. Dimonika, forêt primaire sur crête, sur rocher bien éclairés, 4°13’S, 12°20’E, fr., 02 May 1986, H. de Foresta 983 (P).

### 3. *Ardisia mica* Cheek & Murdoch sp. nov. (Fig. 4)

Syn. *Ardisia sp. no.5* Taton (Taton 1979, p. 120)

**Fig. 4.**
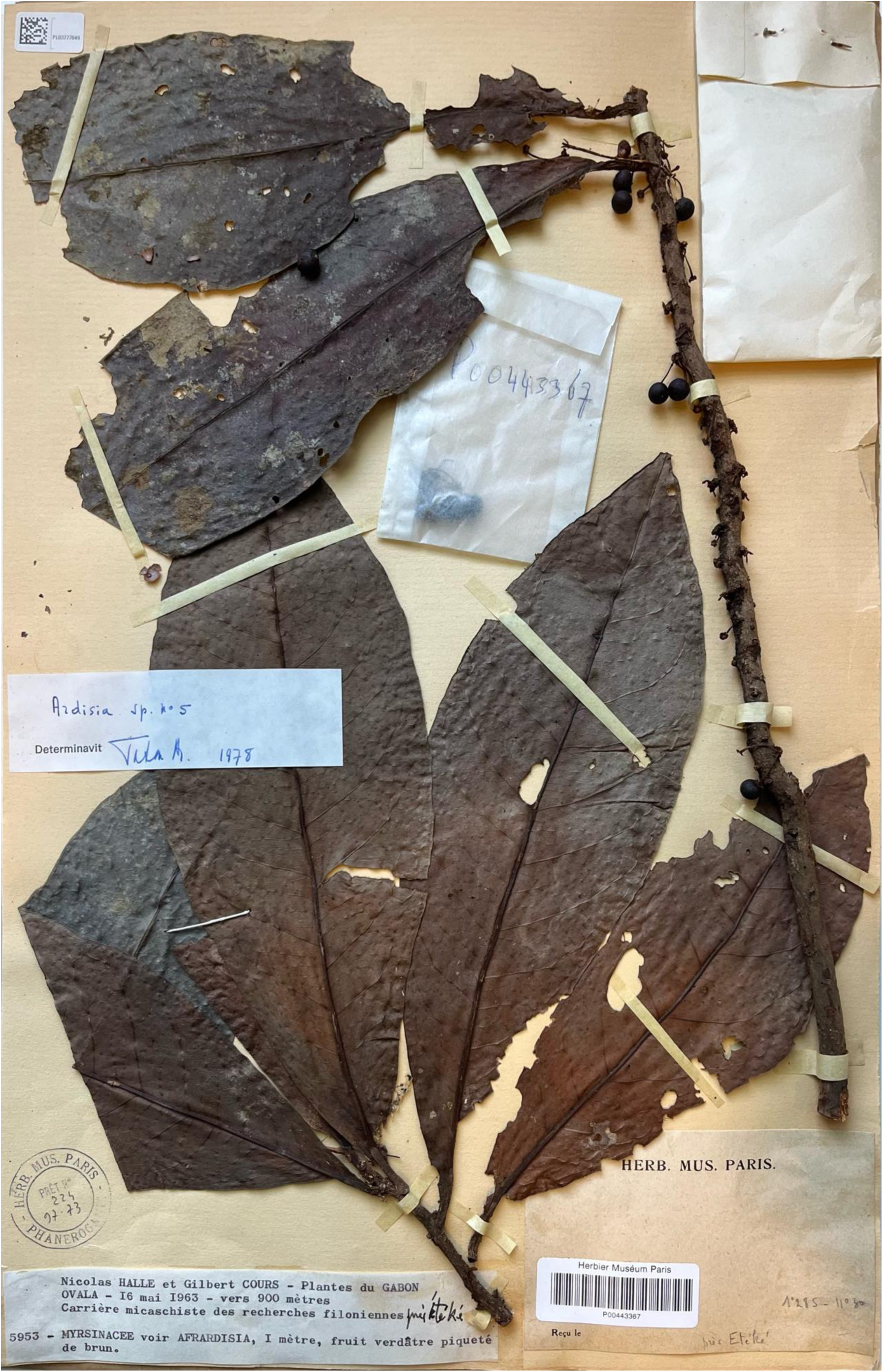
The only known specimen of *A. mica*, *Hallé et Cours* 5953, seen at P.

#### Diagnosis

*Ardisia mica* is most similar to *A. mayumbensis*, differing in stem diameter 6–8 mm vs 3.5–5 mm (at 10 cm above ground); longest leaf-length 24.6 cm vs 18–22 cm; longest leaf width 8.7 cm vs 6.5–7.6 cm; gland density 2–5 per 2 × 2 mm of leaf blade vs 10–27; abaxial leaf scale density 11–14 per 2 × 2 mm vs 18–26. See Table 1 for more diagnostic characters.

#### Type

Gabon, Ngounie, Ovala, Carriere micaschiste des recherches filoniennes, 1°28’S, 11°30’E, 900 m, fr., 19 May 1963, N.Hallé et Cours 5953 (holotype P).

### Etymology

Named as a noun in apposition for the micaschist quarry from where the only known specimen was collected.

### Description

Monocaul, c. 1 m tall, stem vertical, unbranched, phyllotaxy spiral, terete, 6–8 mm diam., internodes 7–17 mm long (excluding the distal four nodes clustered closely together), rusty brown tomentellous, dense in the distal 6 nodes, becoming sparse thereafter, epidermis grey brown to red brown. Leaves clustered towards stem apex, 7 leaves in the distal 10 cm of stem. Petiole canaliculate, 10–12 mm × 2 mm; lamina elliptic, 18–24.6 × 6.6–8.7 cm, length to width ratio 2.8–3:1, decurrent along the petiole at the base, apex acuminate, margin entire, midrib conspicuous, raised on the adaxial and abaxial surfaces, grooved on adaxial; lateral nerves 12–14 on each side of the midrib, departing from midrib at 50–66 degrees, brochidodromous, looping 4–6 mm from margin, tertiary nerves conspicuous, raised abaxially, reticulate creating areolae; oil glands orange-red (transmitted light), brown to black

### Phenology

Fruiting in May, flowering unknown.

### Distribution and ecology

Known from only one collection in Ngounie, central Gabon. 900 m elevation. Ecology unknown.

#### Specimens examined

Gabon. Ngounie, Ovala, Carriere micaschiste des recherches filoniennes, prés Éteké, 1°28’S, 11°30’E, vers 900 m, fr., 19 May 1963, N.Hallé et Cours 5953 (holotype P).

### Conservation status

*Ardisia mica* is known from only one collection in the Ngounie region in central Gabon. A lack of data makes an exact location difficult to pinpoint, but label data tells us that the specimen is collected from within or near a micaschist quarry. Label data also places it close to Etéké Ngounié, now the site of a gold mine (Ebo 2024). The AOO of this species is estimated as 4 km², below the upper threshold for “Critically Endangered” status under Criterion B2. The EOO cannot be calculated due to this species being known from only one occurrence. Given the clear industrial threats to this species, and given it is known from only one collection, this species is provisionally assessed as Critically Endangered B2ab(i,ii,iii,iv,v).

#### Notes

This species is characterised by its elliptic blade, unique among monocaul phanerophyte *Ardisia* of Gabon which generally have more elongated leaves, and being, at c. 1 m tall, the tallest of the species. Its also occurs at a higher altitude than any other species of the group (c. 900 m).

Known from only one specimen, cited by Taton (1979) as *A. sp. No 5*, where he provided a short description and keyed the taxon out, apart from all other known African species of the genus partly on the basis that the flowers are few, and held on the stem below (not among) the leaves. The species is described here in the absence of any other material of the taxon having been found, in order to help improve the conservation outcome of this appraently rare and highly range-restricted taxon. The coordinates cited had been written in pencil on the specimen possibly by Taton. These align broadly with label notes and elevation, that is, close to Eteke and Ovala. We were not able to fully characterise the leaf lamina scale shape due to time constraints at P.

### 4. *Ardisia hallei* Taton (Taton 1979, p. 101). (Fig. 5)

Type: Gabon, Belinga, 1°6’N, 13°14’E, 800 m, b., 31 Oct. 1964, N. Hallé 2953 (holotype P).

**Fig. 5.**
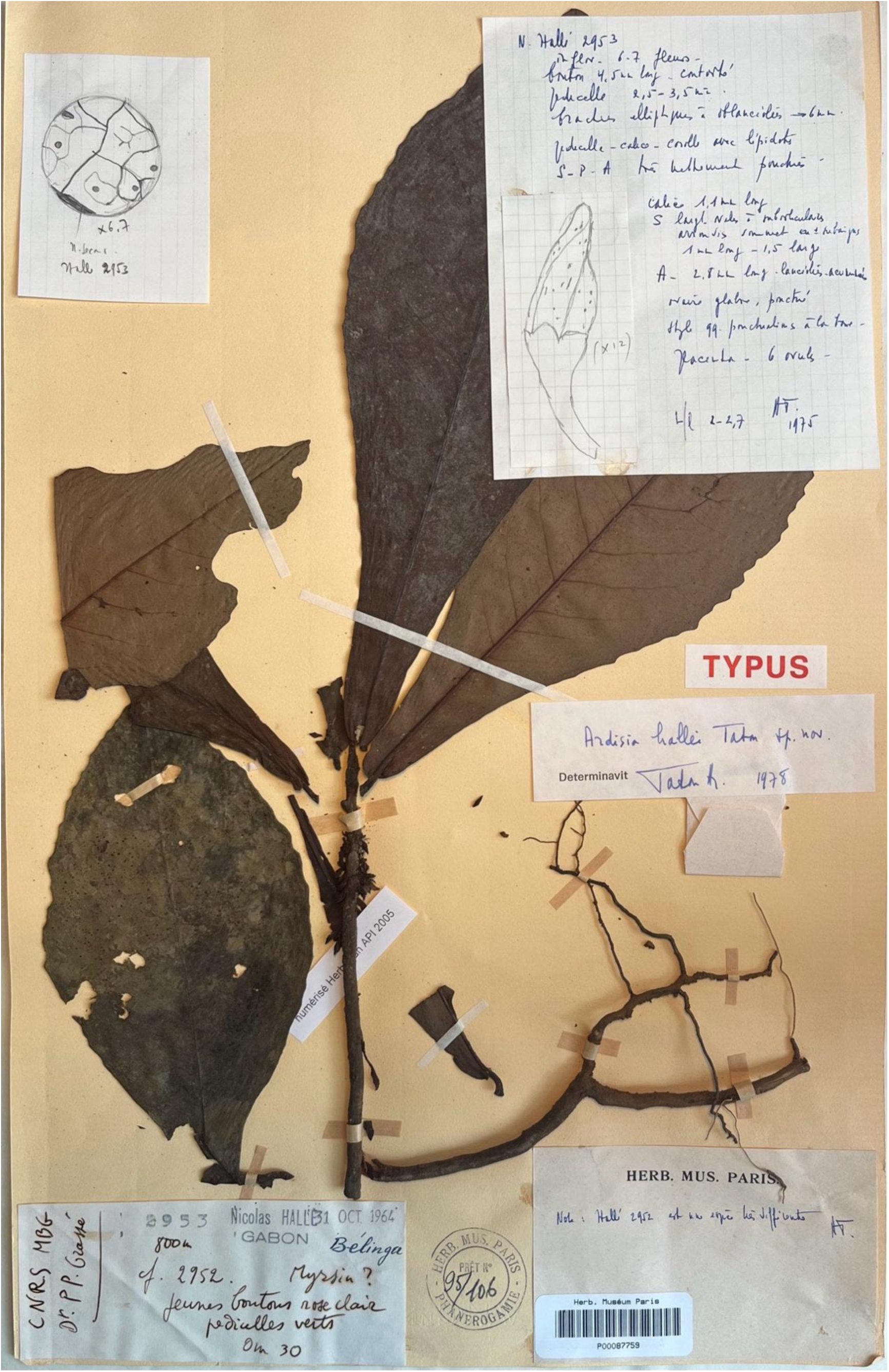
The holotype of *A. hallei*, *Hallé* 2953, seen at P.

#### Etymology

Named for botanist, botanical illustrator, and prolific herbarium specimen collector in Central Africa, Nicolas Hallé, born 1927, who collected the holotype for this species and who was a major contributor to Flore De Gabon.

#### Description

Monocaul, 60–90 cm tall, stem vertical, unbranched, phyllotaxy spiral, terete, 4–7 mm diam.; Adventitious roots absent; Leaves clustered towards stem apex. Petiole canaliculate, 4.5–10 × 2–3 mm; lamina oblanceolate, 14.4–22(23.8) × 5.5–11 cm, length to width ratio 2–2.7:1, apex acute to obtuse, base cuneate then abruptly rounded, margin irregularly crenate, midrib conspicuous, raised on the adaxial and abaxial surfaces, grooved on adaxial; lateral nerves 11–18 on each side of the midrib, departing from midrib at 55–70 degrees, brochidodromous, tertiary nerves conspicuous, raised on both surfaces, reticulate creating areolae; oil glands most dense near margin, subcircular, elliptic or oblong, 0.08–0.7 × 0.08–0.35 mm, 1–3 per 2 mm × 2 mm; 0–6 glands per areole; scales conspicuous (microscope), reddish brown, on abaxial and adaxial surfaces, 9–26 per 2 mm × 2 mm abaxially, 0–8 per 2 mm × 2 mm adaxially. Inflorescences axillary, 2–8-flowered, peduncle usually inconspicuous or up to 1 mm long, rachides 2–3 mm long, pedicel bases exserted from the rachis; bracts elliptic to oblanceolate, 2–5.5 × 1–2.5 mm, oil glands covering 5–10% of surface, dark brown to black; pedicels terete, 2–6(9) × 0.75–1 mm, green in life (N.Hallé 2953). Flower buds narrowly ovoid to narrowly triangular, 3–4 × 1.5–1.75 mm, apex acute, calyx cupular, lobes 5, overlapping to the right, ovate, c. 1.5 × c. 1.5 mm, apex rounded to acute, margin fimbriate, corolla twisted to the right, light pink in life (N.Hallé 2953). Flowers not seen. Immature fruit ovoid, c. 3.25 mm diam., calyx as bud, style base persistent, ovules c. 6, contiguous arrangement. Fruit red in life (J. Florence 1329).

#### Phenology

Flower buds in October, immature fruits in January, June and August.

### Distribution and ecology

Gabon (Belinga) and Republic of the Congo (Sibiti and Kakamoeka). Occurring in lowland to submontane evergreen forest at elevations of 500–870 m.

### Conservation status

*Ardisia hallei* is the only taxon in this treatment to have received a formal IUCN assessment, appearing on the Red List. It is classified as Vulnerable B2ab(i,ii,iii,iv,v) (Paradis et al. 2024). However, this is stated to be on the basis of eight collections at six locations which we suspect may be based on specimens other than this taxon. The photograph used to illustrate this taxon by Paradis et al. (2024) clearly does not represent this species since it shows leaves without the broad marginal teeth and without the strongly tapering and concave proximal leaf-blades of the type.

### Specimens examined

Gabon. Belinga, 1°6’N, 13°14’E, 800 m, b., 31 Oct. 1964, N.Hallé 2953 (holotype P); Belinga, Foret 6 km S du camp. Foret sempervirente, 1°6’N, 13°14’E, 870 m, 13 June 1978, J. Florence 1329 (P). Republic of the Congo. Sibiti, foret pres station. 3°45’S, 13°25’E, 04 Aug. 1965, Farron 4190 (P); Sibiti, Foret sur route Grand-Bois – Madingou, environs de Sibiti, 3°50’S, 13°25’E, 09 Aug. 1965, Farron 4279 (P); Kakamoeka, Route du chantier de Boungolo, foret primaire. Point-Noire. 4°8’S, 12°5’E, 31 Jan. 1966, Farron 4909 (P).

### Notes

*Ardisia hallei* is most similar to *A. mayumbensis*, but is differentiated by its irregularly crenate leaf margin, its much sparser oil glands in the leaf lamina (1–3 per 2 mm × 2 mm vs. 10–27 per 2 mm × 2 mm) and its shorter pedicels (2–6(–9) mm vs (7–)9–15 mm). It is found at elevations of 500–870 m, higher than any known *A. mayumbensis* specimen (10–250 m). Due to limited time at P where the majority of the material is held, this description is less comprehensive than others in this treatment. Some elements of the description are taken from Taton’s protologue (Taton 1979). Fruit not seen in person or digitally, but referred to in field notes of J. Florence 1329.

### 5. *Ardisia bracteata* Baker (Baker in Oliver 1877, p. 495). (Fig. 6)

*Afrardisia bracteata* (Baker) Mez (Mez 1902, p.184). Type as for *Ardisia bracteata* Baker

**Fig. 6.**
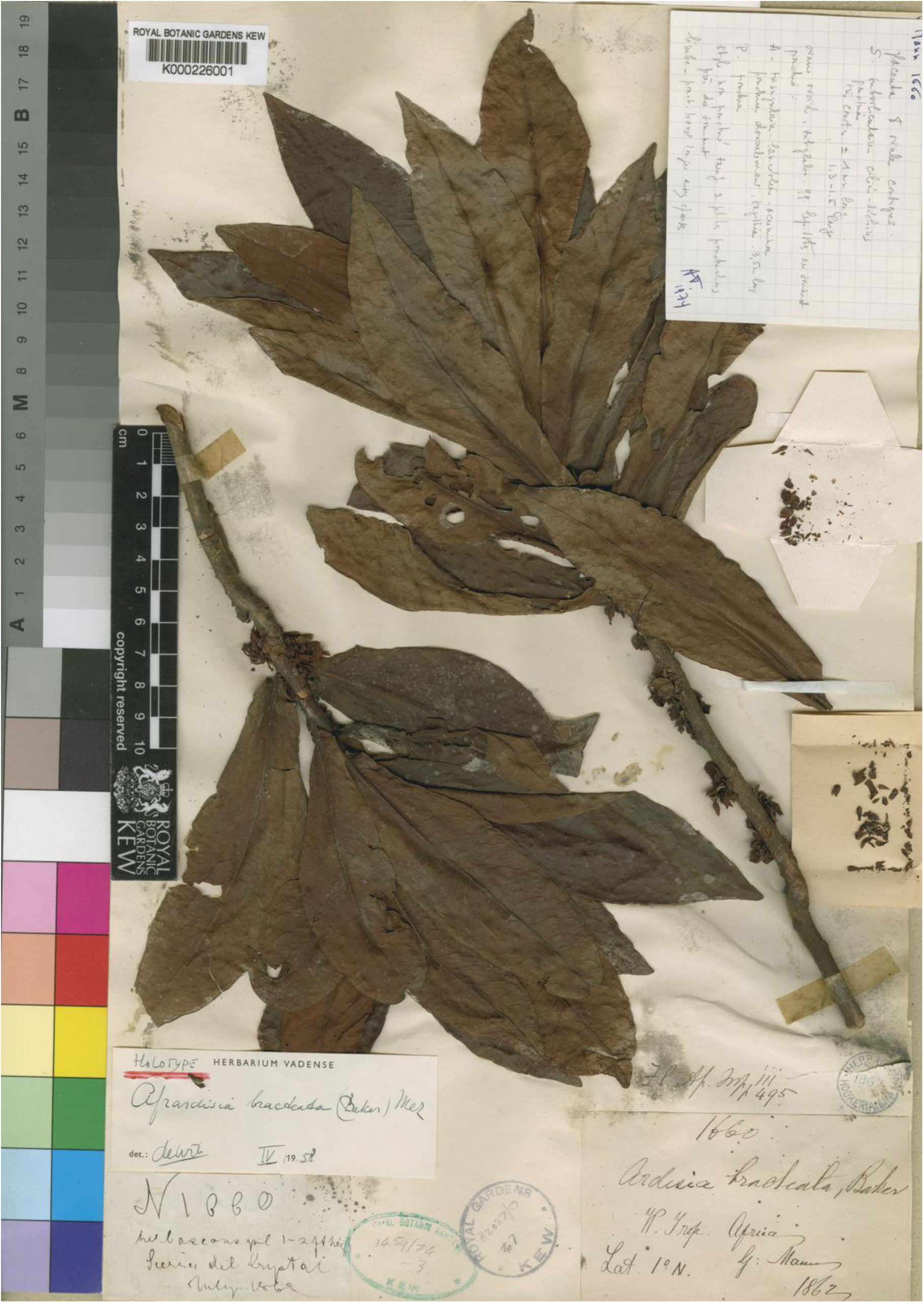
The holotype and currently only known specimen of *A. bracteata*, Mann, *G* 1660, seen at K. (Royal Botanic Gardens, Kew continuously updated).

Type: Gabon, Crystal Mountains, 1°N, b., 1862, Mann, G. 1660 (lectotype K see note below).

#### Etymology

Named for the large bracts found in the inflorescences of this species.

### Description

Monocaul, 30–60 cm tall, stem vertical, unbranched, phyllotaxy spiral, terete, 4.5–6 mm diam., internodes 5–9 mm long; Leaves clustered towards stem apex, 14–17 leaves in the distal c. 5.5 cm of stem. Petiole canaliculate, 3–3.5 × 2–3 mm; lamina oblong-oblanceolate, 8.7–12.8 × 3–3.7 cm, length to width ratio 2.7–3.8:1, apex acute, base narrowing then abruptly rounded, margin subentire, lamina c. 0.8 mm thick, c. 1.2 mm at midrib, midrib conspicuous, raised on the adaxial and abaxial surfaces, grooved on adaxial; lateral nerves 10–16 on each side of the midrib, departing from midrib at 50–75 degrees, brochidodromous, looping 1–1.5 mm from margin, tertiary nerves conspicuous, raised abaxially, reticulate creating areolae; oil glands red-orange (transmitted light), brown to black in reflected light, visible and raised on adaxial and abaxial surfaces in reflected light, densest near margin, subcircular to elliptic, 0.09–0.29 × 0.09–0.24 mm, 2–5 per 2 mm × 2 mm; 0–2 glands per areole; abaxial 2–4 armed scales conspicuous (microscope), reddish brown, c. 23 per 2 mm × 2 mm, 0.06–0.12 mm wide., glabrous adaxially. Inflorescences axillary and cauliflorous, 6–12 flowered, peduncle inconspicuous, rachides 2.5–3 mm long, pedicel bases exserted from the rachis, rusty brown floccose hairs on rachides, bracts and pedicels; bracts obovate, 4.5–10 × 2–6 mm, oil glands covering 5–10 % of surface, subcircular to elliptic, red-brown to black, c. 0.1 × c. 0.1 mm; pedicels terete, 4–7 × 0.66–1 mm. Flower buds narrowly ovoid, 4.5–5 × 1.75–1.25 mm, apex acute; calyx cupular, lobes 5, overlapping to the right, ovate, 0.8–1 × c. 0.8–1.1 mm, apex rounded, oil glands covering 50% of surface, red-brown to black, raised, subcircular to elliptic, c. 0.1 × c. 0.1 mm, margin fimbriate, fimbriae translucent, 0.03–0.1 mm long, corolla twisted to the right. Flowers 5-merous.

Corolla lobes narrowly ovate, c. 4 × 2 mm, apex acute, oil glands covering 5–10% of surface, visible on both sides, red-brown to black, raised, subcircular to elliptic, c. 0.09–0.27 × 0.05– 0.16 mm. Androecium; anthers lanceolate, 3.5–4 × 0.75–1 mm, base subsagittate, apex acute. Ovary subglobose, c. 0.5 mm diam. style terete, c. 3.5 mm long. Fruits not seen

### Distribution and ecology

Known only from the Gabon/Equatorial Guinea border, in the Crystal Mountains. Ecology unknown.

#### Conservation status

*Ardisia bracteata* is known from only one collection close to the border of Gabon and Equatorial Guinea, in the Crystal Mountains. Uncertainty on the precise location of this collection makes it difficult to assess threats. Whether *A. bracteata* still exists within Crystal Mountains National Park, or any other protected area, is unclear. There are at least three logging concessions within the area (World Resources Institute 2014), representing a significant threat to this species. The AOO of this species is estimated as 4 km², below the upper threshold for “Critically Endangered” status under Criterion B2. The EOO cannot be calculated due to this species being known from only one occurrence. Due to the clear threat from logging concessions, as well as the highly restricted AOO, this species is provisionally assessed as Critically Endangered B2ab(i,ii,iii,iv,v).

#### Specimen examined

Gabon. Crystal Mountains, 1°N, b., 1862, Mann, G. 1660 (lectotype K).

### Notes

This species is characterised by the large bracts found in its inflorescences, which are considerably larger than all other monocaul *Ardisia* of Gabon (see Table 1). *Ardisia bracteata* also has the smallest leaves (in terms of both length and width) of all of species of the group, and it lacks the oblong (or dash-like) oil glands in the leaf lamina seen in most African *Ardisia*. However, it does have subcircular and elliptic oil glands.

It is unclear if the specimen is from Gabon or Equatorial Guinea. The supplied coordinates give only “1°N”, placing it directly on the border between the two countries. Here we follow Sosef et al. (2006), which includes it in Gabon. All flowers of the specimen are disassembled, presumably due to dissection by previous workers. We select the K sheet as lectotype in case further duplicates are found, and because it is annotated by the author of the name.

### Key to the monocaul phanerophyte *Ardisia* species of Gabon

1. Adventitious roots present (distal 5–18 nodes) ………………………………..……2 – Adventitious roots absent (distal 5–18 nodes) ……………………………....……..3
2. Pedicels ascending (in fruit), oil glands absent from anthers, gland density (leaf blade, no. per 2 × 2 mm) 2–6, leaf thickness (lamina) 0.7–0.75 mm… 1. *A. doudou* sp. nov. – Pedicels descending (in fruit), oil glands present on anthers, gland density (leaf blade, no. per 2 × 2 mm) 8–16, leaf thickness (lamina) 0.31–0.53 mm …….2. *A. litterbin* sp. nov.
3. Longest leaf length < 24 cm, leaf lamina oblanceolate to oblong ………………… 4– Longest leaf length > 24 cm, leaf lamina elliptic ……………... 3. *A. mica* sp. nov.
4. Longest leaf length >15 cm, width of longest leaf >6 cm, leaf lamina oblanceolate, leaf margin irregularly crenate, largest bracts <6 mm long, bud length (from base of calyx) 3–4 mm, petiole 4.5–10 mm long. ......………...…………………4 4. *A. hallei* – Longest leaf length <13 cm, width of longest leaf c. 3.4 cm, leaf lamina oblong-oblanceolate, leaf margin subentire, largest bracts >6 mm long, bud length (from base of calyx) 4.5–5 mm, petiole 3–3.5 mm long …...…….……...…………5. *A. bracteata*

## Discussion

Among the five monocaul phanerophyte *Ardisia* species of Gabon described here, two are endemic (*Ardisia doudou* and *Ardisia mica*), one is found on the border of Gabon and Equatorial Guinea (*Ardisia bracteata*), and two occur in both Gabon and Republic of the Congo (*Ardisia litterbin* and *Ardisia hallei*). Monocaul phanerophyte species also occur in Cabinda and DRC (*Ardisia mayumbensis*), and additional species occur in Cameroon (species delimitation in progress). All known African monocaul phanerophyte *Ardisia* have a funnel-like cluster of leaves at the stem apex and seem to exhibit some degree of litter-gathering but only two species, *A. doudou* and *A. litterbin* also have adventitious roots in their funnel areas.

### Litter gathering Ardisia

*A. doudou* and *A. litterbin* are unique among all known African *Ardisia* in their use of dense, branching adventitious roots in the apical part of the stem where the funnel of leaves collects falling organic litter. In both species, these roots are among the leaves and, with associated organic matter, can completely conceal the stem surface. These roots are seen in most, but not all specimens, suggesting roots may only develop at certain times. Whether this is seasonal, related to plant maturity, or something else entirely is unclear. It is also possible that roots have been “cleaned” off in the collection process. In herbarium specimens, adventitious roots are brittle and easily broken. Adventitious roots are often densely matted with organic matter. Leaf litter, fungi and invertebrates can be seen. The extent to which these plants may support other lifeforms, potentially in symbiotic relationships, is unknown.

Litter gathering is well established among other angiosperm genera within the same geographic region. Terrestrial litter-gathering species tend to be understory rainforest shrubs with an unbranched stem and a funnel-like rosette of large leaves (Zona and Christenhusz 2015). At least 10 species of West and Central African *Psychotria* (Rubiaceae) exhibit this habit (Lachenaud, 2019). These species are monocaulous or connected by rhizomes, bear large oblanceolate or spathulate leaves at the stem apex and develop adventitious roots on the stem, superficially resembling the litter-gathering *Ardisia* described here. The litterbin *Psychotria* are not monophyletic and are found within several different sections of the genus (Lachenaud and Jongkind 2013). *Calocrater preussii* K.Schum. (Apocynaceae), *Scaphopetalum mannii* Mast. (Malvaceae), *Chassalia ischnophylla* (K.Schum.) Hepper (Rubiaceae), *Coffea magnistipula* Stoff. & Robbr. (Rubiaceae), *Eumachia letouzeyi* (Robbr.) Razafim. & C.M.Taylor (Rubiaceae), *Phyllobotryon spathulatum* Müll.Arg. (Salicaceae), *Allexis cauliflora* (Oliv.) Pierre (Violaceae) are just some of the known terrestrial litter gatherers from Gabon, across several genera and families (Zona and Christenhusz 2015). That this adaptation has evolved many times independently demonstrates its effectiveness. Kenfack et al. (2007) refer to the understorey as being “dominated” by litter-trapping treelets in their census of trees in Korup National Park, southwest Cameroon and litter-gathering species are not infrequent in the lowland forests of that country (e.g. Cheek et al. 2008).

Adventitious roots on the aerial stem near the leaf funnel are an indicative trait of a litter gatherer, but not all litter gatherers have them (Zona and Christenhusz 2015). *Ardisia hallei*, *A. bracteata* and *A. mayumbensis* are referred to by Lachenaud & Jongkind (2013) as litter gatherers. *Ardisia hallei* does not possess adventitious roots but does otherwise resemble a litter gatherer. *Ardisia bracteata* does not possess adventitious roots and has comparatively smaller leaves than any other species in this treatment. Without more material, or observation in the field, it is difficult to say if *A. bracteata* is stically litter-gathering or not. Adventitious roots can be seen in one isotype of *A. mayumbensis* (Gossweiler 7674 at K), but not on isotypes seen digitally at LISC, MO, BM and COI (GBIF.org User 2025a, 2025b). The adventitious roots seen here are branching and in the apical part of the stem, but are sparse and do not contain leaf litter or other organic material as seen in *A. doudou* and *A. litterbin*. It is worth noting that despite having the same collection number, each isotype is a different individual as each includes a stem apex, and monocauls have only a single stem, therefore there is some variation seen among the isotypes of *A. mayumbensis*.

Adventitious roots can also be seen in some creeping, herbaceous African *Ardisia*. These are typically not found at the stem apex but rather on the older stems, rooting them to the ground (Peng and Cheek 2025) and are not connected with a litter-gathering function.

### Conservation of rare *Ardisia* species

While petroleum extraction is currently Gabon’s main source of revenue and does represent a threat to some species in the broader context, Gabon’s oil reserves are running low, and the government are seeking to diversify revenues (International Monetary Fund 2024).

Both sustainable forestry and oil palm plantations are on the agenda (International Monetary Fund (2024). In addition is the $150 million committed to Gabon to protect its forests by the Norwegian government under its Central African Forest Initiative (Mongabay Environmental News 2021). It is unclear at this stage what scale of threat that new forestry and oil palm operations may pose to Gabon’s forests. In our provisional conservation assessments, we have found that all taxa in this treatment fall within threatened IUCN categories, with two, *Ardisia mica* and *A. bracteata*, qualifying as the highest category, Critically Endangered. Logging is a principal threat across these assessments. Regulation of future logging activities will be vital to the conservation of these species as they are each known from so few collections. Logging in Gabon tends to be selective and low density, as it is focussed on one species, *Aucoumea klaineana* Pierre, and is considered by some to be more sustainable and less of a threat to wildlife than elsewhere in Africa (Sullivan et al. 2024). However, its effect on rare plant species in Gabon is unknown. In neighbouring countries such as Cameroon, which has the highest number of recorded plant species extinctions in tropical Africa, oil palm plantations following logging appears to be expanding and are a major concern to rare and threatened plant species survival, while small-holder agriculture, hydro-electric and mining projects are also threats (Murphy et al. 2023). Three out of four new species to science published currently such as those in this paper are threatened with extinction (Brown et al. 2023). Until species have a scientific name it is difficult for them to be accepted onto the IUCN Red List, making it imperative to accelerate the description of species that are still undescribed, since inscription on the Red List can improve the outcome for threatened species (Cheek et al. 2020). Global extinction of plant species is a concern. While some recently described species from Gabon are relatively widespread and may tolerate some human activities (e.g. Cheek et al. 2024), others, known from single collections as is the case with *Ardisia mica* and *A. bracteata*, are feared globally extinct already (Moxon-Holt and Cheek 2011, Cheek et al. 2021). It is notable that these two Ardisia species have not been seen for 164 and 63 years respectively and are also conceivably extinct although further surveys are needed to establish this. Should they be refound we suggest that consideration is given to developing a species conservation action plan for them (e.g. Couch et al. 2022) and that that they be incorporated into designated Important Plant Areas (IPAs) if possible (e.g. as in Murphy et al. (2023) for Cameroon). Although Gabon has designated Key Biodiversity Areas (Texier et al. 2024), these were not designated based on concentrations of threatened plant species or areas of species richness and socio-economic indigenous species as is the case with IPAs (Darbyshire et al. 2017).

### Future research needed

The phylogenetic relationships within African *Ardisia* are poorly known since respectively only four and two African taxa have been included in previous molecular phylogenetic papers (Julius et al. 2012; Larson et al. 2023). Whether the monocaul African *Ardisia* form a distinct clade or not requires much more intensive sampling to resolve. Any future work on this topic should account for the different lifeforms of African *Ardisia* and sample accordingly to investigate if each of these lifeforms forms a monophyletic clade.

This paper, alongside Peng and Cheek (2025), brings further clarity to the our knowledge of the four different lifeforms seen in African *Ardisia* (Cheek *et al*. 2025). However much more basic taxonomic work is needed urgently, first and foremost to delimit and characterise the remaining, shrubby *Ardisia* species which remain unresolved, so that they may be identified, assessed for their conservation status, and where needed, protected.

## Supporting information

Supplemental Table 1 Monocaul Gabon Ardisia

## Notes

### Competing Interest Statement

The authors have declared no competing interest.

## References

Bachman, S., Moat, J., Hill, A., De La Torre, J., and Scott, B. 2011. Supporting Red List Threat Assessments with GeoCAT: Geospatial Conservation Assessment Tool. – ZooKeys 150: 117–26. doi:10.3897/zookeys.150.2109.

Beentje, H. & Cheek, M. 2003. Glossary. In Beentje, (ed.), Flora of Tropical East Africa. – Balkema, Lisse, Netherlands. 115 pp.

Brown, M., Bachman, S., Nic Lughadha, E. 2023. Three in four undescribed plant species are threatened with extinction. – The New Phytologist 240(4): 1340–1344. 10.1111/nph.19214

Cheek, M. and Xanthos, M. 2012. *Ardisia ebo* sp. nov. (Myrsinaceae), a Creeping Forest Subshrub of Cameroon and Gabon. – Kew Bull. 67(2): 281–84. doi:10.1007/s12225-012-9362-8.

Cheek, M., Bissiengou, P. & Lachenaud, O. 2024. *Keetia gordonii* sp. nov. (Rubiaceae - Vanguerieae), a new species of threatened forest liana from Gabon. Kew Bull. 79: 841–853. 10.1007/s12225-024-10219-y

Cheek, M., Horwath, A. and Haynes, D. 2008. *Psychotria kupensis* (Rubiaceae) a new dwarf, litter-gathering species from western Cameroon. – Kew Bull. 63: 243–246. 10.1007/s12225-008-9018-x

Cheek, M, Harvey, Y. and Onana, J. 2010. The Plants of Dom, Bamenda Highlands, Cameroon: A Conservation Checklist. – Royal Botanic Gardens, Kew, UK.

Cheek, M., McDaniel, C. and Murdoch, H. 2025. Classification of Life Forms of African *Ardisia* (Myrsinaceae or Primulacae) (Abstract Only). – AETFAT Bulletin 50: 23.

Cheek, M., Nic Lughadha, E., Kirk, P., Lindon, H., Carretero, J., Looney, B., Douglas, B., Haelewaters, D., Gaya, E., Llewellyn, T., Ainsworth, M., Gafforov, Y., Hyde, K., Crous, P., Hughes, M., Walker, B.E., Forzza, R.C., Wong, K.M., Niskanen, T. 2020. New scientific discoveries: plants and fungi. – Plants, People Planet 2: 371–388. 10.1002/ppp3.10148

Cheek, M., Tchiengué, B., van der Burgt, X. 2021. Taxonomic revision of the threatened African genus *Pseudohydrosme* Engl. (Araceae), with *P. ebo*, a new, critically endangered species from Ebo, Cameroon. PeerJ 9 :e10689 10.7717/peerj.10689

Couch, C., Molmou, D., Magassouba, S., Doumbouya, S., Diawara, M., Diallo, M. Y., Keita S.M., Koné, F., Diallo, M.C., Kourouma, S., Diallo, M.B., Keita, M.S., Oularé, A., Darbyshire, I., Gosline, G., Nic Lughadha, E., van der Burgt, X, Larridon, I. & Cheek, M. (2022). Piloting development of species conservation action plans in Guinea. Oryx, 1–10. 10.1017/s0030605322000138

Darbyshire, I., Anderson, S., Asatryan, A., Byfield, A., Cheek, M., Clubbe, C., Ghrabi, Z., Harris, T., Heatubun, C. D., Kalema, J., Magassouba, S., McCarthy, B., Milliken, W., Montmollin, B. de, Nic Lughadha, E., Onana, J. M., Saıdou, D., Sarbu, A., Shrestha, K & Radford, E. A. 2017. Important Plant Areas: revised selection criteria for a global approach to plant conservation. Biodivers. Conserv. 26: 1767 – 1800. 10.1007/s10531-017-1336-6.

De Wit, H. C. D. 1958. Revision of *Afrardisia* Mez (Myrsinaceae). – Blumea 4 (supplement): 242–262

Ebo, P. 2024. Etéké Ngounié: Uni Mine d’or Dans Les Mains Du Marocain Managem. 2024. – TV Plus Afrique. https://www.tvplusafrique.com/index.php?page=detail-article&article=4112&titre=ETÉKÉ NGOUNIÉ : UNE MINE D’OR DANS LES MAINS DU MAROCAIN MANAGEM (accessed: August 6, 2025).

GBIF.org User. 2025a. Occurrence Download. – doi:10.15468/DL.UYJY3J.

GBIF.org User. 2025b. Occurrence Download. – doi:10.15468/DL.UYQNB2.

Gilg, E. 1902. Myrsinaceae africanae. – Botanische Jahrbücher Für Systematik Pflanzengeschichte Und Pflanzengeographie. 30: 96–101.

Gilg, E. and Schellenberg, G. 1913. Myrsinaceae africanae. II. – Botanische Jahrbücher Fur Systematik, Pflanzengeschichte Und Pflanzengeographie. 48: 512–525.

Good, R.D. 1927. Myrsinaceae. – Journal of Botany, British and Foreign 65 (supplement): 69.

Google Earth Pro. Continuously updated. Satellite Imagery of Gabon. – https://www.google.com/earth/ (July 21, 2025).

International Monetary Fund 2024. Gabon’s Diversification Journey: What is Missing? International Monetary Fund Africa Dept. – https://www.elibrary.imf.org/view/journals/002/2024/145/article-A003-en.xml

IUCN 2012. IUCN Red List Categories and Criteria: Version 3.1. Second Edition. – International Union for Conservation of Nature, Gland and Cambridge. http://www.iucnredlist.org/.

IUCN Standards and Petitions Committee. 2024. Guidelines for Using the IUCN Red List Categories and Criteria. Version 16. – Prepared by the Standards and Petitions Committee. https://www.iucnredlist.org/documents/RedListGuidelines.pdf.

Julius, A., Gutiérrez-Ortega, J. S., Sabran, S., Tagane, S., Naiki, A., Darnaedi, D., … and Kajita, T. 2021. Phylogenetic relationship of tropical Asian *Ardisia* and relatives (Primulaceae) shows non-monophyly of recognized genera and subgenera. – Shokubutsu Kenkyu Zasshi (Journal of Japanese Botany) 93: 149–165.

Kenfack, D., Thomas, D.W., Chuyong, G., Condit, R. 2007. Rarity and Abundance in a Diverse African Forest. – Biodiversity and Conservation 16(7): 2045–74. doi:10.1007/s10531-006-9065-2.

Lachenaud, O. (2019). Révision du genre Psychotria (Rubiaceae) en Afrique occidentale et centrale. Opera Botanica Belgica 17. Jardin botanique de Meise. 909 pp.

Lachenaud, O. and Jongkind, C.C.H. 2013. New and Little-Known *Psychotria* (Rubiaceae) from West Africa, and Notes on Litter-Gathering Angiosperms. – Plant Ecology and Evolution 146(2): 219–33. doi:10.5091/plecevo.2013.765.

Larson, D., Chanderbali, A., Maurin, O., Gonçalves, D.J.P., Dick, C., Soltis, D., Soltis, P. et al. 2023. The Phylogeny and Global Biogeography of Primulaceae Based on High-Throughput DNA Sequence Data. – Molecular Phylogenetics and Evolution 182: 107702. doi:10.1016/j.ympev.2023.107702.

Leaf Architecture Working Group. 1999. Manual of Leaf Architecture: Morphological Description and Categorization of Dicotyledonous and Net-Veined Monocotyledonous Angiosperms. – Washington, DC:

Massicotte, P. and South, A. 2025. Rnaturalearth: World Map Data from Natural Earth. – doi:10.32614/CRAN.package.rnaturalearth.

Mez, C.C.M. 1902. Myrsinaceae. – Das Pflanzenreich :Regni Vegetablilis Conspectus. 9(4): 183–87.

Mishra, Shivam. 2024. Gabon’s National Oil Company to Buy Assala Energy from Carlyle. – Offshore Technology. https://www.offshore-technology.com/news/gabon-to-buy-assala-energy/ (August 6, 2025).

Mongabay Environmental News. 2021. Gabon Becomes First African Country to Get Paid for Protecting Its Forests. – https://news.mongabay.com/2021/07/gabon-becomes-first-african-country-to-get-paid-for-protecting-its-forests/ (August 6, 2025).

Moxon-Holt, L. and Cheek, M. 2021. *Pseudohydrosme bogneri* sp. nov. (Araceae), a spectacular Critically Endangered (Possibly Extinct) species from Gabon, long confused with *Anchomanes nigritianus*. – Aroideana 44(1): 110–131. 10.1101/2021.03.25.437040

Murphy, B., Onana, J.M. van der Burgt, X. M., Tchatchouang Ngansop, E., Williams, J., Tchiengué, B., Cheek, M. 2023. Important Plant Areas of Cameroon. – Royal Botanic Gardens, Kew. https://kew.iro.bl.uk/concern/books/c056b5cb-b146-4509-b98b-a7f5dd49517e

Oliver, D. 1877. Mysinaceae. – Flora of Tropical Africa. 3: 495–96. London, L. Reeve and Co.

Onana, J.-M. and Cheek, M. 2011. Red Data Book of the Flowering Plants of Cameroon: IUCN Global Assessments. – RBG, Kew. 578pp.

Paradis, A.-H., Bidault, E. and Stévart, T. 2024. Ardisia hallei. – The IUCN Red List of Threatened Species 2024: E.T207931284A207934300. doi:10.2305/IUCN.UK.2024-1.RLTS.T207931284A207934300.en.

Pebesma, E. 2018. Simple Features for R: Standardized Support for Spatial Vector Data. – The R Journal 10(1): 439–46. doi:10.32614/RJ-2018-009.

Pebesma, E. and Bivand, R. 2023. Spatial Data Science: With Applications in R. – Chapman and Hall/CRC. doi:10.1201/9780429459016.

Peng, P. and Cheek M. 2025. A synoptic revision of the creeping herbaceous African species of *Ardisia* (Primulaceae or Myrsinaceae) with six new species from Cameroon and Gabon. ––Webbia 80(2) Suppl.: 139–168. doi: 10.36253/jopt-19150

POWO continuously updated. Plants of the World Online. – Facilitated by the Royal Botanic Gardens, Kew. Published on the Internet; https://powo.science.kew.org/ Retrieved 04 April 2025.

R Core Team. 2025. R: A Language and Environment for Statistical Computing. – Vienna, Austria: R Foundation for Statistical Computing. https://www.R-project.org/.

Royal Botanic Gardens, Kew continuously updated. The Herbarium Catalogue. – http://www.kew.org/herbcat (June 10, 2025).

Sosef, M.S.M., Wieringa, J.J., Jongkind, C.C.H., Achoundong, G., Azizet Issembé, Y., Bedigian, D., Van Den Berg, R.G., Breteler, F.J., Cheek, M., Degreef, J. 2006. Checklist of Gabonese Vascular Plants. – Scripta Botanica Belgica 35. National Botanic Garden of Belgium. 435 pp.

South, A., Schramm, M., and Massicotte, P. 2024. Rnaturalearthdata: World Vector Map Data from Natural Earth Used in ‘Rnaturalearth’. – doi:10.32614/CRAN.package.rnaturalearthdata.

Sullivan, M. K., Vleminckx, J., Bissiemou, P. A. M., Niangadouma, R., Mayoungou, M. I., Temba, J. L., … & Comita, L. S. (2024). Low-intensity logging alters species and functional composition, but does not negatively impact key ecosystem services in a Central African tropical forest. – Global Ecology and Conservation 53 e02996. 10.1016/j.gecco.2024.e02996

Swartz, O. 1788. Nova Genera & Species Plantarum; Seu, Prodromus Descriptionum Vegetabilium, Maximam Partem Incognitorum. Holmiae [etc.] – In Bibliopoliis Acad. M. Swederi. doi:10.5962/bhl.title.4400.

Taton, A. 1979. Contribution a l’etude du genre *Ardisia* Sw. (Myrsinaceae) en Afrique tropicale. – Bulletin du Jardin botanique national de Belgique / Bulletin van de National Plantentuin van België 49(1/2): 81. doi:10.2307/3667819.

Texier, N., Ngama, S., Essomba, G. N., and Stévart, T. 2024. Les Zones Clés pour la Biodiversité du Gabon. – Missouri Botanical Garden & Université Libre de Bruxelles, Brussels, 220 pp.

UNEP-WCMC and IUCN. 2025. Protected Planet: The World Database on Protected Areas (WDPA) and World Database on Other Effective Area-Based Conservation Measures (WD-OECM). – http://protectedplanet.net/ (August 14, 2025).

Wickham, H. 2016. Ggplot2: Elegant Graphics for Data Analysis. – Springer-Verlag New York. https://ggplot2.tidyverse.org.

Wickham, H., François, R., Henry, L., Müller, K. and Vaughan, D. 2023. Dplyr: A Grammar of Data Manipulation. – doi:10.32614/CRAN.package.dplyr.

Wickham, H., Hester, J. and Bryan, J. 2024. Readr: Read Rectangular Text Data. – doi:10.32614/CRAN.package.readr.

De Wildeman, E., and Bequaert, J.C. 1925. 3 Plantae Bequaertianae: Études Sur Les Récoltes Botaniques Du Dr. J. – Bequaert, Chargé de Missions Au Congo Belge (1913–1915). Gand: A. Hoste. https://catalog.hathitrust.org/Record/002001005.

World Resources Institute. 2014. Global Forest Watch. – www.globalforestwatch.org. (July 31, 2025).

Yang, C. and Hu, J. 2022. Molecular Phylogeny of Asian *Ardisia* (Myrsinoideae, Primulaceae) and Their Leaf-Nodulated Endosymbionts, *Burkholderia* s.l. (Burkholderiaceae). – PLOS ONE 17(1): e0261188. doi:10.1371/journal.pone.0261188.

Zona, S., and Christenhusz, M.J.M. 2015. Litter-Trapping Plants: Filter-Feeders of the Plant Kingdom. – Bot. J. Linn. Soc. 179(4): 554–86. doi:10.1111/boj.12346

