## Supplemental Table 1 Monocaul Gabon Ardisia for "A Taxonomic Revision of the Monocaul Phanerophyte *Ardisia* (Primulaceae) of Gabon"

Table 1. Diagnostic characters separating the monocaul *Ardisia* species of Gabon, Republic of the Congo and Cabinda. Longest leaves are the longest measurable leaf of each collection. *A. mayumbensis* measurements taken from *Gossweiler* 7674 [K!], *Nkondi, F.* 673 [K!] and *J. Hombert* 450 [K!].

|  | *Ardisia doudou* | *Ardisia litterbin* | *Ardisia mica* | *Ardisia hallei* | *Ardisia bracteata* | *Ardisia mayumbensis* |
| --- | --- | --- | --- | --- | --- | --- |
| Elevation (m) | 170-450 m | 10-460 m | 900 m | 500 – 870 m | Unknown | 10-250 m |
| Range | Doudou Mountains (Gabon) | Western Gabon, Mayombe Forest (Republic of the Congo) | Ngounie (Central Gabon) | Belinga (Gabon) & SW Republic of the Congo | Crystal Mountains (Gabon/Equatorial Guinea border) | Mayombe Forest (Cabinda, Republic of the Congo, DRC) |
| Height | 50-60 cm | 30-70 cm | 100 cm | 60 – 90 cm | 30-60 cm | 60-90 cm |
| Stem diameter at 10cm above ground | 5-11 mm | 6-8 mm | 6-8 mm | 4-7 mm | 4.5-6 mm | 3.5-5 mm |
| Leaf shape | Oblanceolate | Oblanceolate | Elliptic | Oblanceolate | Oblong-oblanceolate | Oblanceolate to elliptic |
| Longest leaf, length to width ratio | 2.4-3.4:1 | 2.4-3.6:1 | 2.8:1 | 2-2.8:1 | 3.8:1 | 2.7-3.1:1 |
| Longest leaf, length range | 28.3-33.3 cm | 23.2-30 cm | 24.6 cm | 15.8-22(23.8) cm | 12.8 cm | 18-22 cm |
| Longest leaf, width range | 8.6-14 cm | 7.7-12 cm | 8.7 cm | 6-11 cm | 3.4 cm | 6.5-7.6 cm |
| Leaf thickness (lamina) | 0.7-0.75 mm | 0.31-0.53 mm | Unknown | Unknown | c. 0.83 mm | 0.25-0.31 mm |
| Pedicel posture | Ascending (in fruit) | Descending | Descending | Descending | Descending | Descending |
| Gland density (leaf blade, no. per 2mm x 2 mm) | 2-6 | 8-16 | 2-5 | 1-3 | 2-5 | 10-27 |
| Glands per areole (leaf blade reticulation) | 1-4(6) | 5-35 | 1-3 | 0-6 | 0-2 | 4-55 |
| Scale density (leaf blade, abaxial surface, no. per 2 mm x 2 mm) | 14-22 | 14-23 | 11-14 | 9-26 | c. 23 | 18-26 |
| Adventitious roots amongst the distal stem nodes | Usually present | Usually present | Absent | Absent | Absent | Rarely present |
| Bracts size | 2-5 x 0.7-1.2 mm | 2-6 x 0.6-3 mm | 2-4 x 1-2 mm | 2-5.5 x 1-2.5 mm | 4.5-10 x 2-6 mm | 2-4 x 0.8-1.6 mm |
| Petiole length | 4-13 mm | 5-16 mm | 10-12 mm | 4.5-10 mm | 3-3.5 mm | 7-20 mm |
